# Brain dynamics of classical psychedelics show paradoxical hierarchical flattening with increased complexity

**DOI:** 10.1101/2024.12.21.629922

**Authors:** Jakub Vohryzek, Elvira Garcia-Guzmán, Morten L. Kringelbach, Edmundo Lopez-Sola, Christopher Timmermann, Leor Roseman, Enzo Tagliazucchi, Giulio Ruffini, Robin Carhart-Harris, Gustavo Deco, Yonatan Sanz Perl

**Affiliations:** Center for Brain and Cognition, Computational Neuroscience Group, Department of Information and Communication Technologies, Universitat Pompeu Fabra, Barcelona, Spain; Centre for Eudaimonia and Human Flourishing, Linacre College, University of Oxford, UK; Department of Psychiatry, University of Oxford, Oxford, United Kingdom; Centre for Music in the Brain, Aarhus University, Aarhus, Denmark; Brain Modeling Department, Neuroelectrics, 08035 Barcelona, Spain; Center for Psychedelic Research, Department of Brain Sciences, Imperial College London, London SW7 2AZ, UK; Universidad de Buenos Aires, Facultad de Ciencias Exactas y Naturales, Departamento de Física, and CONICET - Universidad de Buenos Aires, Instituto de Física Aplicada e Interdisciplinaria (INFINA). Buenos Aires, Argentina; Latin American Brain Health Institute (BrainLat), Universidad Adolfo Ibañez, Santiago, Chile; Departments of Neurology and Psychiatry, University of California San Francisco, San Francisco 94143, USA; Institució Catalana de la Recerca i Estudis Avançats (ICREA), Barcelona, Spain; Departamento de Matemática y Ciencias, Universidad de San Andrés, Buenos Aires B1644BID, Argentina; National Scientific and Technical Research Council (CONICET), Ciudad Autónoma de Buenos Aires C1425FQB, Argentina

## Abstract

Despite divergent behavioral and phenomenological profiles, both psychedelic states and reduced states of consciousness have been associated with a flattening of the brain’s functional hierarchy. To address this apparent paradox, we developed a more specific definition of hierarchy based on the proximity of the brain to thermodynamic equilibrium and then applied it to investigate the changes to the functional hierarchy elicited by three classical serotonergic psychedelics – psilocybin, lysergic acid diethylamide, and dimethyltryptamine. We found that all three psychedelics consistently induced a global reduction in the functional hierarchy. In contrast to the flattening of the functional hierarchy observed during loss of consciousness, psychedelics displaced the brain towards equilibrium while simultaneously increasing the complexity of neural activity, indicating a unique mechanism linked to specific changes in the configuration and differentiation of resting-state networks. This work showcases how metrics based on statistical mechanics can be used for the specific characterization of different global brain states, contributing to the understanding of consciousness as a collective process emerging from complex neural interactions.

## Introduction

Psychedelics have garnered substantial attention in recent years for their ability to induce a profound, acute, and long-term functional reorganization within the human brain, with concomitant changes to behavior, cognition, and subjective experience^1^. Understanding these changes is important not only for a comprehensive grasp of the underlying neurobiology but also for discovering the mechanisms supporting potential therapeutic applications^2–4^. A fruitful approach to address these questions has focused on the functional interactions between brain regions, establishing that psychedelic action induces a reduction in the brain’s functional hierarchy^5–9^. Nevertheless, this reduction has also been observed in other conditions that are behaviorally different from the acute effects of psychedelics, such as deep sedation, general anesthesia, and disorders of consciousness^10,11^, thus severely limiting the interpretation of these findings as specific signatures of altered states of consciousness.

These apparent contradictions might stem from the lack of precise definitions of what constitutes a ‘hierarchy’ in neuroscience, with multiple definitions coexisting in the literature^12^. In general, hierarchy in biological systems can be mathematically described using order theory and it plays a fundamental role in living systems. However, neuroscience has yet to apply rigorous hierarchical analysis, as noted by Hilgetag and Goulas in a recent perspective^13^. In particular, brain functional hierarchy can be best understood through measures capturing segregation versus integration, reflecting local versus global network dynamics. The concept of a “global workspace” suggests that a small group of brain regions integrate and broadcast information, orchestrating hierarchical function, akin to a conductor managing an orchestra^14^. In that direction, transfer entropy, a method for measuring information flow, has been used to identify global workspace regions^15^. Alternative approaches have implemented connectivity gradients to describe functional hierarchical organization^16^. Recently, an emerging alternative is a thermodynamics-based framework, which offers a more efficient way to quantify hierarchical organization and can help uncover the brain’s generative mechanisms^17^.

Regarding the study of brain organisation under psychedelics, recent literature has shown that psychedelic interventions desynchronise the brain^1^ and change its organisation by shifting the spatial differentiation of brain functional connectivity. Unimodal regions, related to sensory cortices, become less constrained from the transmodal regions, related to higher-level cognitive cortices; leading to novel perceptual experiences^18^. In that direction, recent evidence has shown that the principal gradient representing such functional organisation is compressed under psilocybin, lysergic acid diethylamide (LSD), and dimethyltryptamine (DMT)^5,8^. Furthermore, harmonic decomposition of functional interactions has been shown to collapse under DMT-induced brain state^7^. Overall, different approaches to investigating brain organisation provide clear evidence of the flattening of the functional hierarchy of the brain during psychedelics.

In this work, we investigated the hierarchical organisation during psychedelics based on a precise definition of the brain’s hierarchical organization using thermodynamics principles to quantify the temporal asymmetry in information flow, termed Directed Functional Hierarchy (DFH). This theoretical framework defines the ‘breaking of the detailed balance’ as a direct measure of asymmetry for any physical system and determines the level of non-equilibrium^17,19–21^. On the one hand, recent studies have shown higher levels of temporal asymmetry (i.e. more nonequilibrium) during cognitive tasks, implying a more hierarchical organization needed for the specific computations^19,22,23^. On the other hand, a lower level of irreversibility (i.e. less nonequilibrium) has been observed in disorder-of-consciousness human patients, anesthetized non-human primates, and in a deep sleep of ferrets suggesting constrained computational capabilities^19,24,25^. Additionally, investigating the network hierarchical organisation has revealed differential ways of how psilocybin and escitalopram (a first-line antidepressant treatment) work in rebalancing brain dynamics in depression long-term post-intervention^26^, has implicated the importance of fronto-striatal-thalamic circuitry in long-term effects of psilocybin^27^, and has shown a decrease in higher temporal frequencies under LSD as measured by magnetoencephalography^28^.

Here, we assessed the effects of psychedelics on the brain’s DFH and neural complexity by leveraging previously published datasets of resting-state functional magnetic imaging (fMRI) from healthy human participants under the effects of three drugs, namely psilocybin^29^ (9 participants), LSD^30^ (15 participants) and DMT^8^ (17 participants), all in the acute post-intervention stage. Based on the fact that the non-equilibrium nature of a system is inherently tied to its spatiotemporal resolution^31^, we also investigated whether changes under psychedelic drugs are consistent across different spatial scales by varying the graining of the parcellation from 100 to 1000 brain regions in the Schaefer parcellations^32^. To further complement the mechanisms driving the psychedelic state we studied the level of complexity of the signals and its relationship to the level of hierarchy. Finally, we computed the network and spatial differentiation of DFH to study the functional consequences of the changes to the DFH changes.

We hypothesized that the acute effects of psychedelics would induce a global reduction of DFH in comparison to control participants. As the DFH in brain dynamics is intrinsically related to the spatial scale, we further conjecture that there is an optimal parcellation for describing the psychedelic effects. In contrast to the changes observed in states of reduced consciousness, we hypothesized increased brain complexity alongside the flattening of the functional hierarchy, indicative of different mechanisms implicated in both sets of conditions. Finally, we expected to find a re-organization at the level of resting-state networks in parallel to the observed changes to emergent collective dynamics, consistent with previous reports of psychedelic action on the architecture of large-scale brain activity.

## Results

### Overview

Our approach is based on a principled measure of hierarchy inspired by the theory of thermodynamics, introducing the DFH as the amount of asymmetry in the dynamics of the brain. We quantified the extent of temporal asymmetry in terms of time-shifted correlations, computed as the pairwise Pearson’s correlation of the time series with a temporal shift between pairs of signals. In that way, we obtained an asymmetrical matrix that contains how the signal *j* in time *t* + Δ*t* is affected by the signal *i* in time *t* in the entry *ij*, and vice versa in the entry *ji* (**see Methods**). We then measured global DFH by computing the Fano factor over the distribution of pairwise values of the magnitude of temporal asymmetry between various regions (**Figure 1B**) and computed DFH for three different psychedelic-induced brain states – DMT, LSD, and Psilocybin (**Figure 1A and C**). We also studied the impact of coarse-graining across 6 different parcellations (**Figure 1D**). Furthermore, we investigated the relationship between the DFH under psychedelics and Brain Complexity, as measured by global LZ complexity (**Figure 1E**). Finally, we analysed the functional consequences of the reduction in global DFH at the resting-state networks level calculating within and between network DFH, and in terms of functional differentiation of the DFH matrix for the two principal components (**Figure 1F**).

**Figure 1:**
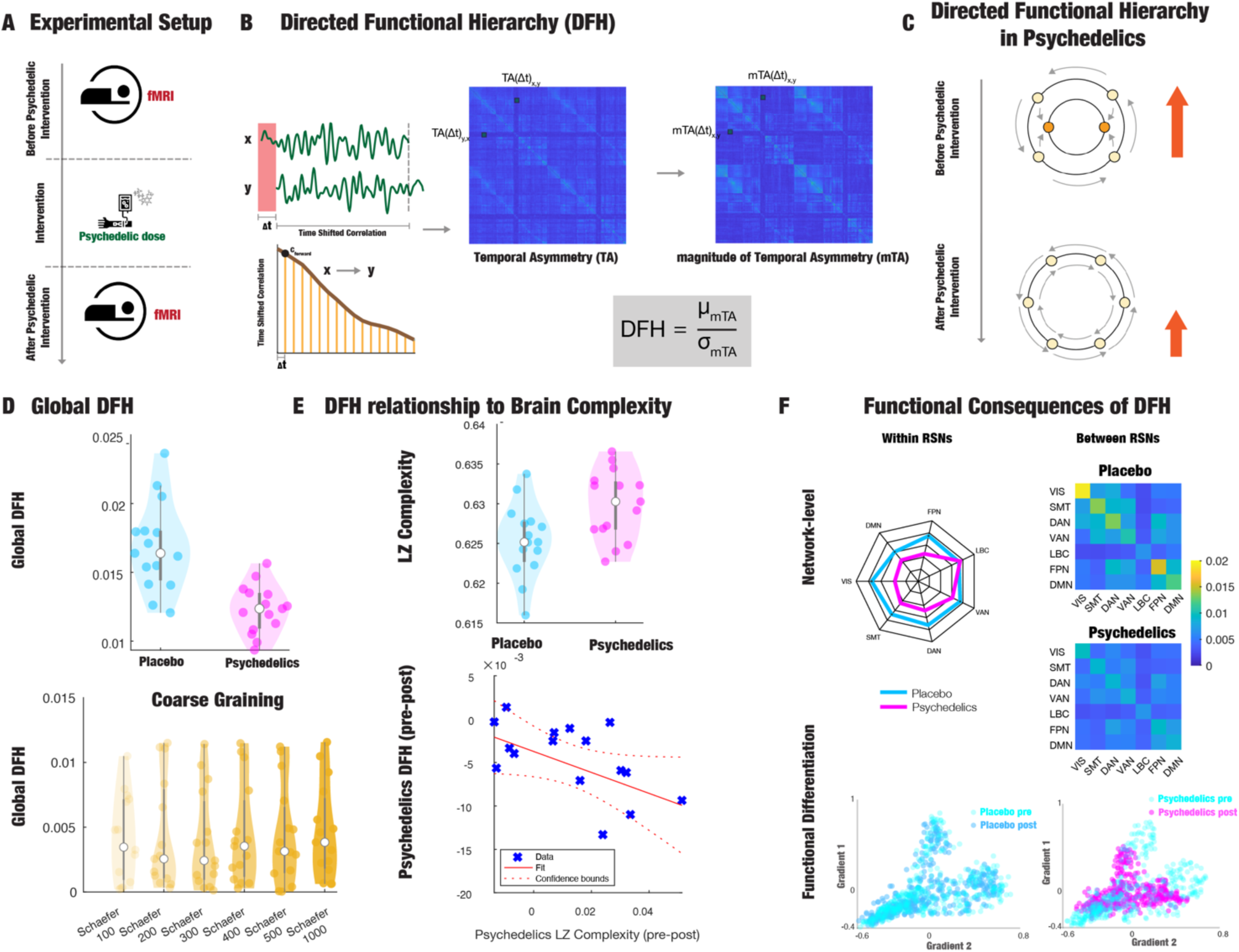
Calculation of Brain Directed Functional Hierarchy in the Acute Psychedelic State. **A)** Neuroimaging fMRI data of whole-brain dynamics before and during the intervention with psychedelic compounds – psilocybin (n=9), LSD (n=15), and DMT (n=17) (see Methods). **B)** The pairwise temporal asymmetry is measured as a time-shifted correlation between brain regions of interest. At the whole-brain level, the temporal asymmetry is summarized in an asymmetric matrix represented by the Time Shifted Correlation matrix. The Magnitude of temporal asymmetry is defined as the square of the difference between time-shifted correlations of regions of interest. **C)** The working hypothesis is that directed functional hierarchy (DFH) in the psychedelic-induced state decreases compared to the pre-intervention state. **D)** Global DFH is calculated as the Fano Factor of the distribution of regional magnitudes of temporal asymmetry. The influence of coarse-graining on the Global temporal asymmetry was performed for different spatial scales defined by the multiscale Schaefer parcellation. **E)** We evaluated the link between global DFH and Brain Complexity (LZ Complexity) pre- and post-psychedelics intervention. **F)** Functional consequences to the changes of global DFH at the network level and in terms of functional differentiation, derived for the two principal components of the DFH matrix.

### Global Directed Functional Hierarchy is flattened during psychedelics states

We investigated how the DFH is reorganized during a psychedelic state based on a thermodynamics definition capturing the directionality of temporal information flow. We computed the temporal asymmetry from the fMRI time series extracted from different psychedelic drugs and the corresponding placebo condition (**Figure 2)**. We then define the temporal asymmetry matrix as the absolute value of the difference between the shifted correlation matrix and its transpose (a symmetric square matrix). We quantified the global level of the magnitude of temporal asymmetry by computing the Fano factor, i.e., the ratio between the mean and the standard deviation across all entries of these matrices, for each participant in each condition and dataset.

**Figure 2.**
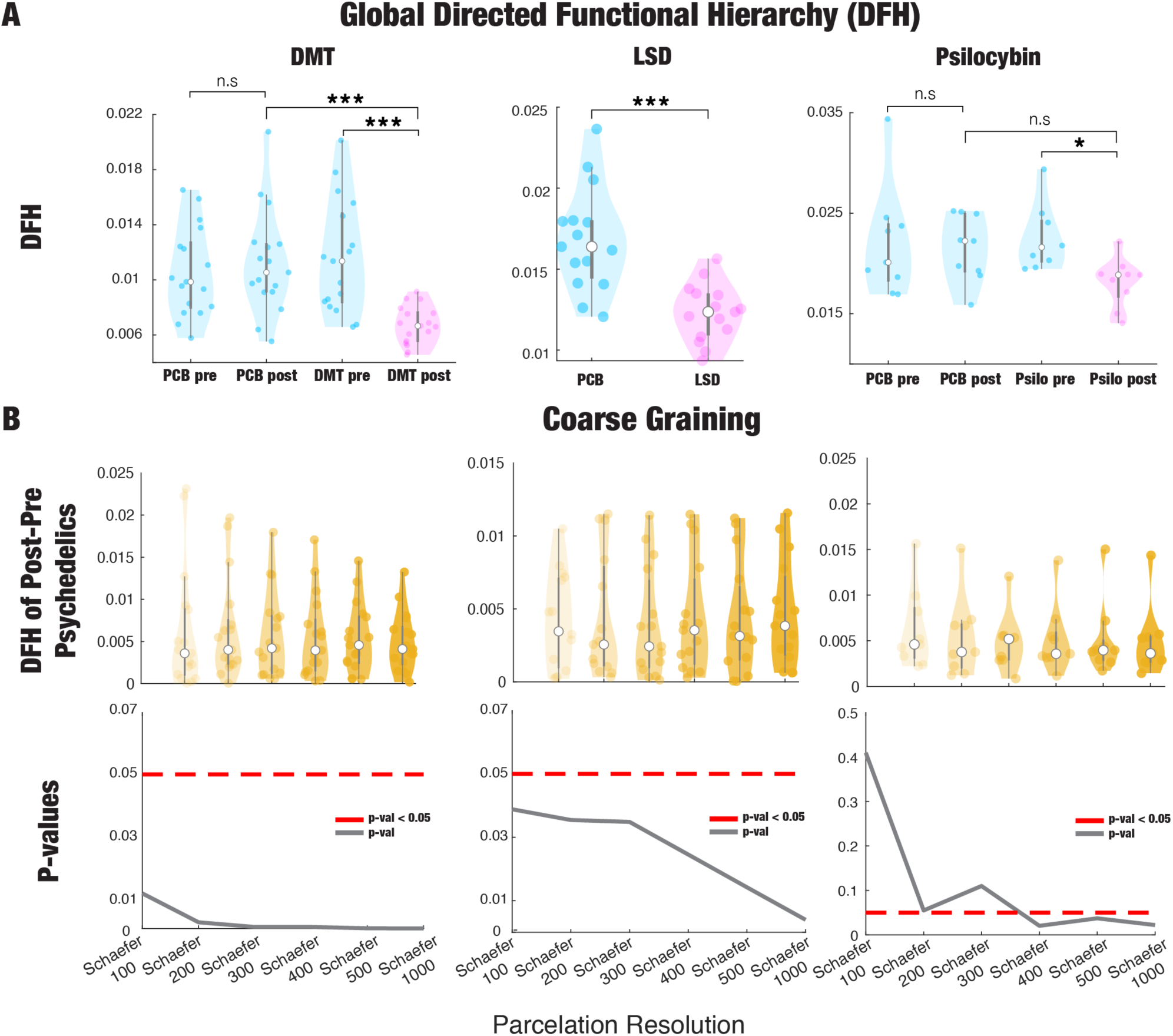
Global reduction of Directed Functional Hierarchy during psychedelic effects. **A)** We computed the DFH for each condition in the three psychedelic datasets for Schaefer 1000 parcellation. We found that in the post-psychedelic administration condition (pink violin plots) the DFH is significantly lower compared with the placebo condition (PCB) pre and post-injection and compared with the before-drug administration (blue violin plots) (two-tailed paired t-test, * p<0.05, *** p<0.001). **B)** We found that the differences between before and after dose administration increases with the resolution of the parcellation while becoming more significant in terms of the reduction of the p-value (two-tailed paired t-test) stabilizing around Schaefer 400 parcellation.

We found that the global DFH significantly decreases under the effects caused by different psychedelic drugs (DMT (DMT post vs. DMT pre, p<1×10^-4^, DMT post vs. PCB post, p<1×10^-4^, LSD (LSD vs. PCB, p=3×10^-4^) and psilocybin (PSILO post vs. PSILO pre, p=0.022, PSILO post vs. PCB post, p=0.057) (**Figure 2A**). However, we did not find differences before and after the placebo (DMT: PCB post vs. PCB pre, p=0.419, PSILO: PCB post vs. PCB pre, p=0.79). These results were obtained using a fine-grained parcellation consisting of 1000 cortical brain regions^32^. Based on these changes, we can interpret that the directed functional hierarchy is flattened during the psychedelic state.

Considering that the DFH of a system is dependent on the spatial scale, we also investigated the effect of spatial graining through different-sized cortical parcellations on the hierarchical reorganization during the psychedelic state. To do so, we exhaustively computed the DFH for six different brain parcellations ranging from 100 to 1000 regions as defined by Schaefer and colleagues^32^. In **Figure 2B** we show how the difference between the DFH before and after psychedelic states increases with finer parcellations, while the statistical comparison becomes more significant, as measured by the reduction of the p-value of a paired t-test. Importantly, we noticed that for the three drugs differences between pre and post-dose administration are significant in all the cases for parcellation resolutions of more than 300 brain regions. For the DMT and LSD drugs, the results are significant across all parcellation resolutions.

### Brain Complexity is anticorrelated to the reduction of DFH during psychedelic states

In order to understand how the reduction in DFH in an acute psychedelic state differs from the similar reduction in hierarchy under reduced states of consciousness such as deep sedation, coma, and disorder of consciousness, we first investigated the level of global brain complexity, measured by the mean of LZ complexity across all brain regions. Furthermore, to understand the possible mechanisms driving the reduction in DFH we correlated the changes in global DFH to the changes in global brain complexity.

For the post-psychedelic administration condition, the global LZ Complexity was significantly higher compared with the post placebo condition for all three psychedelics, and compared with the before-drug administration *DMT* (two-tailed paired t-test, DMT post vs. DMT pre, p<0.012, DMT post vs. PCB post, p<0.005, LSD (LSD vs. PCB, p=0.041) and psilocybin (PSILO post vs. PSILO pre, p=0.210, PSILO post vs. PCB post, p=0.023) (**Figure 3A**). However, we did not find differences before and after the placebo (DMT: PCB post vs. PCB pre, p=0.279, PSILO: PCB post vs. PCB pre, p=0.125). Furthermore, we found that the intervention difference in global LZ Complexity was anti-correlated with the global DFH. This is statistically significant for both DMT and Psilocybin but not LSD (Spearman Correlation DMT: R = -0.52, p=0.034, Psilocybin: R = -0.77, p=0.021, LSD: R = - 0.11, p=0.704) (**Figure 3B**). We calculated the global LZ Complexity in the Schaefer 400 parcellation, which is the resolution with stable results in reshaping the DFH during psychedelics (see **Figure 2**).

**Figure 3.**
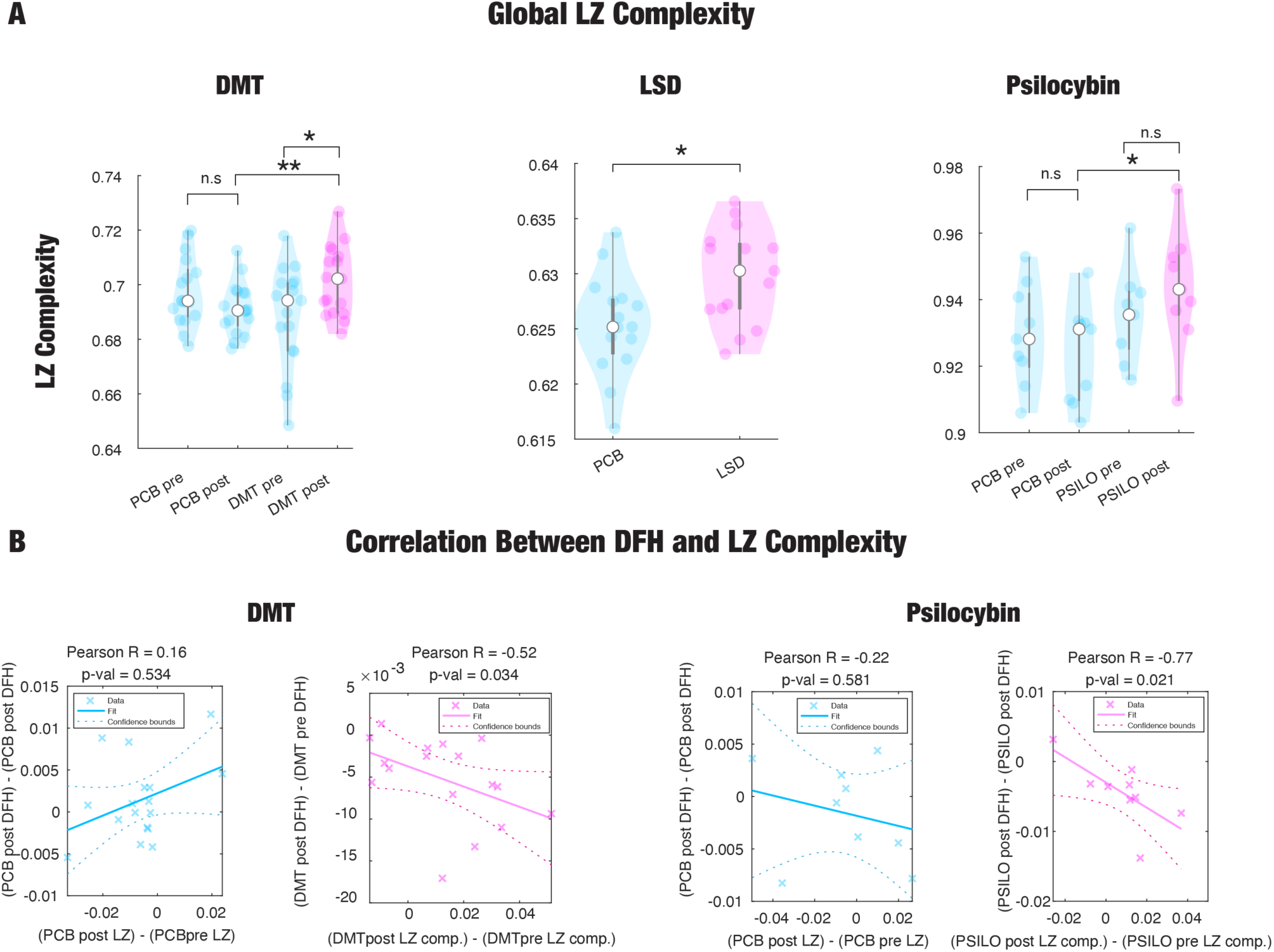
Brain Complexity is anticorrelated to the reduction of DFH during psychedelic states. **A)** The Global LZ Complexity for each condition in the three psychedelic datasets for the Schaefer 400 parcellation, which is the resolution with stable results in reshaping the DFH during psychedelics (**see** Figure 2). We found that post-psychedelic administration condition (pink violin plots) the global LZ Complexity was significantly higher compared with the placebo condition (PCB) pre and post-injection and compared with the before-drug administration (blue violin plots) (two-tailed paired t-test, * p<0.05, ** p<0.01). **B)** Pre and post-psychedelic intervention differences in global LZ Complexity are anticorrelated with the global DFH. This is statistically significant for both DMT and Psilocybin.

### Network-level DFH is reconfigured during psychedelic states

Following our results at the global level, we focused here on the changes that DFH undergoes at the functional system level, reflecting the role that different networks play in hierarchical reconfiguration. We computed the DFH for each of the functional resting state networks (RSN), namely the Yeo 7 RSN^33^, for each participant with each psychedelic drug and the placebo condition (PCB) before and after administration. The within RSN DFH was computed by averaging weights within a network, and between RSN DFH was computed by averaging weights between two networks. First, we investigated how the DFH is modified within each RSN across conditions (**Figure 4A**). We found reduced DFH for the default mode network (DMN), frontoparietal network (FPA), and dorsal attention networks (DAN) for the three psychedelics, (DMT: DMT post vs. DMT pre; VIS - p<10^-4^, SMT - p<10^-4^, DAN – p=0.014. VAN - p=0.035. LBC p=0.809, FPA - p<10^-4^, DMN - p=0.0052, PCB post vs. PCB pre; n.s., LSD: LSD vs. PCB; VIS - p<10^-4^, SMT – p=0.054, DAN - p=0.014. VAN - p=0.015. LBC p=0.184, FPA – p=0.036, DMN – p=0.002, PSILO: PSILO post vs. PSILO pre; VIS – p=0.335, SMT – p=0.474, DAN – p=0.012. VAN – p=0.851. LBC p=0.448, FPA – p=0.015, DMN – p=0.006, PCB post vs. PCB pre; n.s). Secondly, we assessed the reconfiguration of DFH interactions between RSNs during the psychedelic state induced by different drugs. In **Figure 4B**, we display the interaction matrices before and after dose administration (DMT and psilocybin) and after dose administration and placebo condition (DMT, LSD, and psilocybin). We found that interactions involving the DMN and FPA networks were significantly altered during the psychedelic state compared with the PCB condition and before dose administration **(Table S1)**. Importantly, for the DMT and psilocybin datasets, we noticed almost no significant differences when comparing before and after PCB administration, supporting the hypothesis that psychedelic drugs are responsible for reshaping DFH interactions.

**Figure 4.**
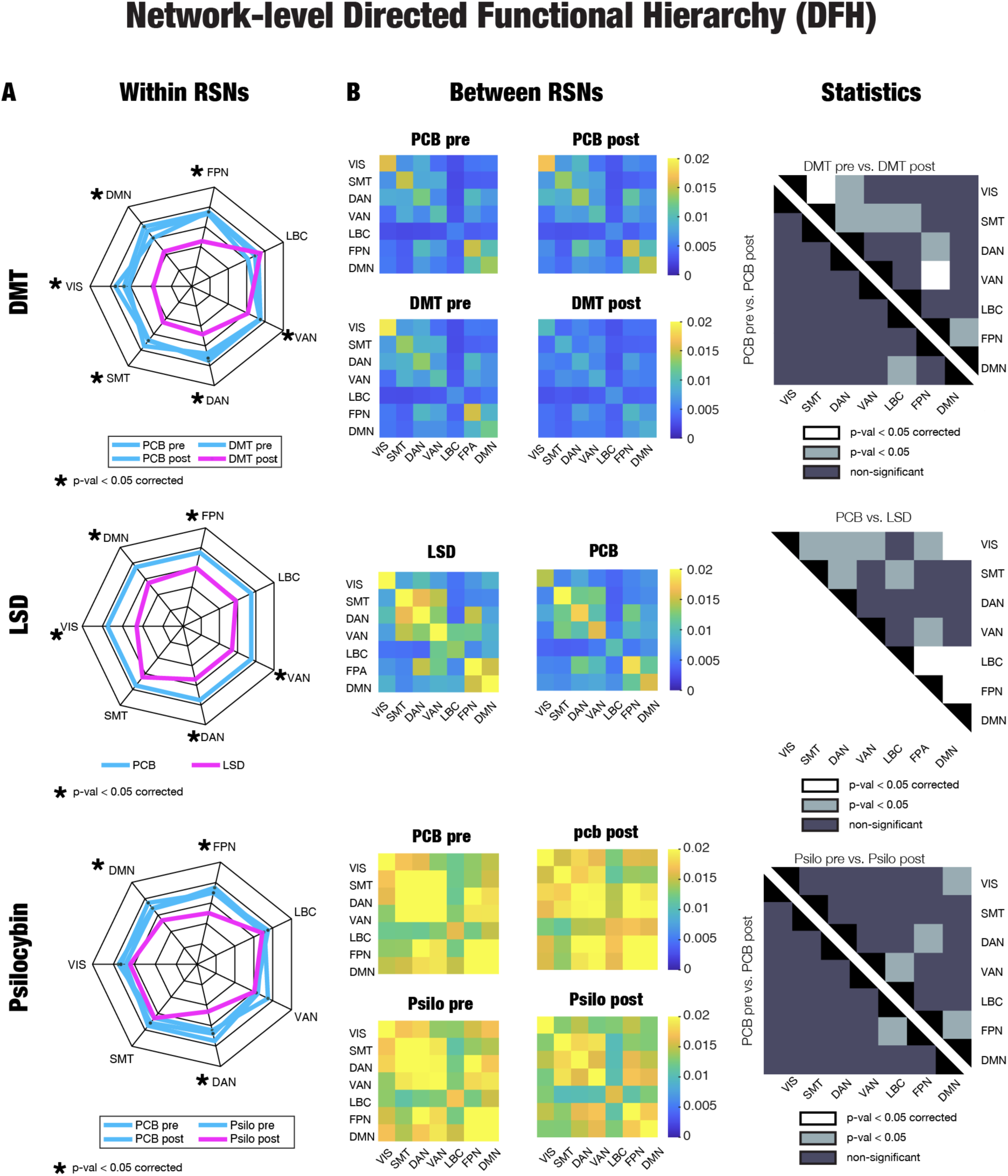
Functional system Directed Functional Hierarchy is reconfigured during psychedelic states. We computed the DFH for each functional system, namely the Yeo 7 resting state networks, and for each condition in the three datasets. **A)** We investigated how the DFH of each RSN is altered during the psychedelic state compared with the PCB condition and before the dose administration. We found that DMN, FPA, and DAN significantly decrease DFH during psychedelic states. In the figure, we report fdr corrected p-values (* p<0.05 corrected). **B)** The interaction matrices for each dataset and each condition averaged across participants (first column). We assessed the statistical differences between conditions in terms of paired t-test false discovery rate corrected (white entries) and without correction (grey entries) (rightmost column) (see Table S1). (DMN: Default Mode; FPA: Frontoparietal; LBC: Limbic; VAN: Ventral Attention; DAN: Dorsal Attention; SMT: Somatomotor; VIS: Visual networks).

### Functional differentiation of DFH is reshaped during psychedelics

Another perspective on the consequences of reduced DFH under psychedelics pertains to the brain’s functional differentiation. Typically, functional differentiation is assessed using principal component analysis or similar nonlinear methods (Diffusion Map Embeddings, Laplacian Eigenmodes, among others ^34^) to extract the components from functional data. These methods provide a low-dimensional representation of the data by computing the most significant patterns of variation of functional connectivity (FC) in the form of gradients. Specifically, the first functional gradient captures the greatest variation in the functional connectivity organisation^35^. In previous studies, functional gradients were examined during the acute phase following psychedelic administration, revealing a collapse of the first gradient during the DMT state measured as the difference between the minimum and maximum range of the functional data projected in that first gradient^8^. Here, we hypothesize that the functional differentiation of the DFH will also be reshaped during the effects of psychedelics, showing differences in the range of the projections of DFH on the principal components. While typically the number of components is selected arbitrarily, often focusing on the first gradient because of its primary importance as it explains most of the variance in the data, here we chose the number of components that have given us the same variance explained across each dataset.

For each dataset, we obtained the first two components of DFH, as we found that this is the minimum number of components showing no significant differences in the explained variance (**Figures S1 and S2**). Consequently, for the rest of the components analysis, we considered the first two components to assess how functional differentiation of the DFH is reshaped under psychedelics. **Figure 5A** shows the spatial distribution of the first two DFH components for the three different psychedelics drugs in Schaefer 400 parcellations using PCA methods implemented in the BrainSpace Toolbox^34^ with regional associations to each RSN for each condition in each dataset. For all three datasets, we noticed that the first components separated the Somatomotor network from Default Mode and Frontoparietal networks, and the second gradient separated the Visual from higher-order networks. At the individual level, we investigated the contraction of the gradients in the two-dimensional space determined by the first two components. We projected the loading of each brain region in this two-dimensional space for each drug and we quantified the components contraction by computing the surface spanned by the loading projections in each case. We found that these components were lower during psychedelic state (two-tailed paired t-test, DMT dataset: DMT post vs. DMT pre, p=3×10^-4^, DMT post vs. PCB post, p=5×10^-4^, PCB post vs. PCB pre, p=0.594, LSD dataset: LSD vs. PCB, p=0.007, PSILOCYBIN dataset: PSILO post vs. PSILO pre, p=0.026, PSILO post vs. PCB post, p=0.883, PCB post vs. PCB pre, p=0.357), showing that psychedelics induce a two-dimensional components contraction of the DFH functional organization (**Figure 5B**).

**Figure 5.**
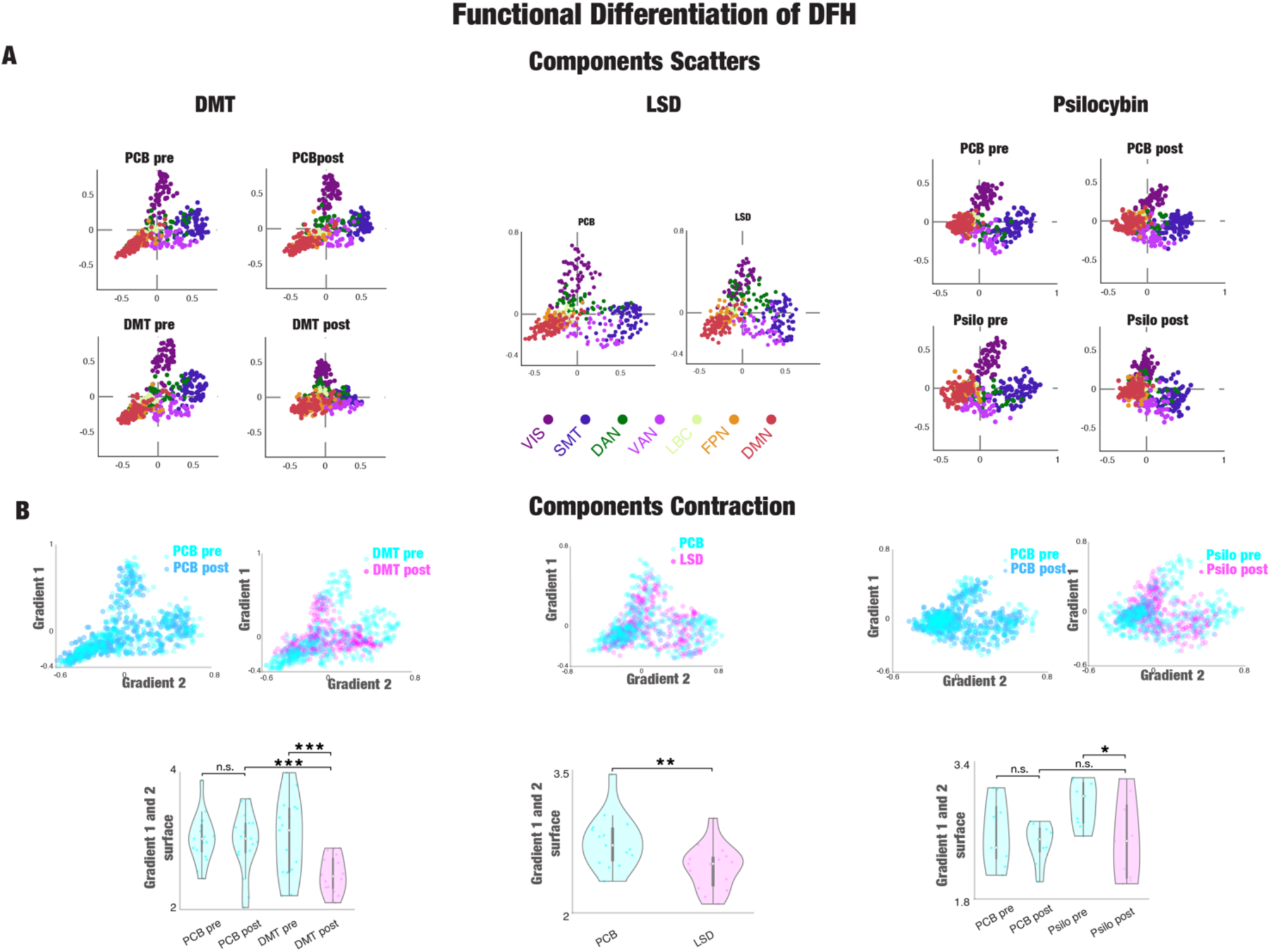
Functional differentiation of DFH is reconfigured under a psychedelic state. **A)** We obtain the first two functional components of the DFH for the three datasets (DMT, LSD, and PSILOCYBIN). The timeseries for this analysis were extracted in Schaefer 400 parcellation, which is the minimum resolution showing significant results in reshaping the DFH during psychedelics (**see** Figure 2). In the three cases, the spatial distribution of both components is similar for the three drugs. We found that the amount of variance explained is not significantly different for the three datasets pre and post-drug, and PCB administration when the first two components are considered (see **Figure S1**). We related the components loading, i.e., the projection of each brain region into the components, with each RSN for each condition in each dataset for the first two components. Consistently for the three datasets, the first gradient separates the Somatomotor network with Default Mode and Frontoparietal networks, and the second gradient separates the Visual with higher-order networks. **B)** We quantified the reshaping of the functional organisation of the DFH by computing the surface spanned by the projections of the loadings in the two-dimensional space generated by the first two components. We found a gradient surface contraction during psychedelic states (two-tailed paired t-test, * p<0.05, ** p<0.01, *** p<0.001).

## Discussion

We introduced a precise definition of the brain’s functional hierarchy in terms of the temporal asymmetry of information flow between brain regions and used it to investigate the effect of psychedelics on large-scale brain activity. Our findings suggest that the acute effects of psychedelics lead to a collapse of the Directed Functional Hierarchy at the global level, compared with control conditions. Furthermore, we demonstrate that spatial graining impacts the estimation of DFH during the psychedelic state, highlighting the importance of the appropriate spatial scale in deriving the measure of DFH. Together with the increase in global brain complexity and the anticorrelation between global DFH and complexity, our results distinguish the psychedelic state from states of reduced states of consciousness such as deep sedation, coma, and disorders of consciousness, also suggesting that psychedelics displace brain activity closer to the thermodynamic equilibrium through a different mechanism that those indexed solely by the increase in global brain complexity. Our analysis also revealed that the transmodal networks show a consistent decrease in the reconfiguration of DFH under all the studied serotonergic psychedelics. Moreover, the analysis of the spatial differentiation of DFH indicates a collapse of the first two principal components during the acute psychedelic effects. Importantly, these results imply that under psychedelics the brain’s dynamics are driven closer to the thermodynamic equilibrium through a mechanism that simultaneously increases global brain complexity, in contrast to what is observed in states of reduced consciousness. We posit that the observed changes in the DFH imply a flattening of the functional hierarchy underlying spontaneous neural activity, enabling a more flexible and accessible brain orchestration.

### A thermodynamically grounded framework to study the functional hierarchy and its application to psychedelics

The notion of functional hierarchy is not precisely defined in neuroscience^17^, mainly due to the different perspectives and methods targeting specific aspects of brain organisation^12^. In this paper, we applied a precise definition of the hierarchy of a system, namely Directed Functional Hierarchy, that computes the level of temporal asymmetry in the directionality of information flow^17^. This description stems from the theory of thermodynamics which posits that the ‘breaking of the detailed balance’ serves as a direct measure of asymmetry for any physical system. Indeed, asymmetry is a key feature in recurrent neural networks as well as in the human brain because it allows for directed information flow, dynamic feedback, and the emergence of complex temporal behaviors which are crucial for computation^36^. As an example, asymmetrically connected neuronal populations enable the network to maintain and evolve internal states over time, a key requirement for tasks involving memory and sequence prediction, among other aspects of computation^37^. In contrast, symmetric or bidirectional connections often cancel out these dynamic effects, limiting the network’s ability to compute complex temporal patterns effectively^38–41^. Such a definition of Directed Functional Hierarchy based on temporal asymmetry differs from previously used approaches that expressed the flattening of functional hierarchies under psychedelics in terms of compression of the spatial differentiation derived from static functional connectivity^5,8^. Intuitively this can be understood by appreciating that the Directed Functional Hierarchy encodes the amount of directionality of information flow, which is not possible to obtain in traditional approaches that use static and undirected functional connectivity.

Psychedelic action enhances the richness of spatiotemporal dynamics by desynchronizing the brain and disrupting functional connectivity^1^. This has been supported by a broadening of the repertoire of functional states and an increase in temporal complexity^42–46^, corroborating theoretical and empirical works positing that psychedelics collapse the brain’s functional hierarchies, “flattening the landscape” of brain dynamics^18,47,48^. By escaping deep local minima under psychedelics, the brain practically fosters flexible and adaptative patterns of thought and behavior^49^. Here, the lower levels of global and network Directed Functional Hierarchy compared to control conditions align with these findings (**Figures 2 and 4**). Nevertheless, because our measure of hierarchy is based on thermodynamics, the insights we gain from our results differ from those of previous undirected approaches which do not capture the asymmetry of the information flow between brain regions.

Previous theoretical work has proposed that the increase of the brain’s complexity under psychedelics indexes the richness of subjective experience^50–52^. However, it is not clear a priori if this change relates to shifts towards or away from thermodynamic equilibrium. Consistent with our results, related work on directed functional connectivity in higher frequencies of MEG data under psychedelics observed a breakdown of information flow^28,53^. Notably, we have found a global decrease in the Directed Functional Hierarchy under all three psychedelics compared with the control condition (**Figure 2**). As expected, we observed no statistical differences in the global DFH before and after the placebo intervention. In this light, the reduction of DFH under psychedelics can be related to the known level of flattening of spatially differentiated functional networks^5,8,54^.

Studies on undirected functional connectivity have also shown that the influence of psychedelics is exerted differently across the brain. Psychedelics tend to decrease within network connectivity while increasing between network connectivity^55,56^. This is further supported by the components of functional differentiation showing their contraction during psychedelic action. Our results also show a different spatial influence of psychedelics as calculated for the seven canonical resting-state networks^57^ (**Figure 4**). The temporal asymmetry is consistently reduced across the three psychedelic drugs within the Default Mode Network, Frontoparietal Network, and Dorsal Attention Network. Interestingly, these networks belong to the transmodal end of the principal gradient of functional organisation^35^. Similarly, the same trend of reduced DFH is found in the interactions across networks that specifically involve DMN and FPN. On the other hand, the unimodal networks, such as Visual and Somatomor networks, also exhibit a reduced DFH in the interactions between networks although they are not statistically significant after FDR correction. The decreased temporal asymmetry within and between RSNs demonstrates an altered computational capacity under the effects of psychedelics which are mainly driven by transmodal networks.

Inspired by the work of Margulies et al.^16^ we assessed the spatial differentiation of the DFH. We found two principal components of DFH (**Figure 5**) explaining the equal amount of variance for each condition (**Figures S1 and 2**). The first component separates the SMN from DMN and FPN while the second component separates the VIS from the higher-order networks (DMN, FPN, VAN) (**Figure S3**). Indeed, these two components of temporal asymmetry show a two-dimensional contraction under the effects of psychedelics. Independently of the specific networks, all regions show a trajectory of moving away from the components’ extremes. This, in turn, reflects a flattening of the spatial differentiation which probably captures both within and between network reduction in temporal asymmetry. All in all, the reductions in DFH support the notion of brain dynamics being closer to the thermodynamic equilibrium.

### Thermodynamic implications of the collapse of the Directed Functional Hierarchy

To support life, physical systems must operate far from thermodynamic equilibrium. Langton’s seminal work also showed that maximal computational capability emerges in dynamical systems poised at the “edge of chaos,” a regime between order and randomness where complex, structured patterns and universal computation can arise^58^. This insight established a link between criticality and information processing, suggesting that non-equilibrium conditions are integral to meaningful computation. More recently, England’s theoretical advances demonstrated that life-like behavior and adaptive organization can naturally emerge as dissipative structures under sustained energy flow^59,60^. By framing biological systems as inherently far-from-equilibrium entities capable of “computing” probabilistic inferences about their environment, this perspective aligns with the notion that non-equilibrium conditions are not only conducive to complexity but essential for sustaining processes that both generate and utilize information in a functionally meaningful way. Landauer’s principle further provides a thermodynamic foundation for this relationship, explicitly linking information processing to energy dissipation and entropy production^61^. This connection extends to Friston’s free energy principle^62^, which conceptualizes biological systems as nonequilibrium steady-state engines that minimize variational free energy—a statistical and information-theoretic quantity—while continuously dissipating energy to maintain organized, inference-driven states. Together, these lines of inquiry unify thermodynamics, information processing, and adaptive computation under the umbrella of nonequilibrium physics for the study of living systems.

As mentioned above, systems performing computations must operate out of equilibrium, as computation inherently involves information processing, state changes, and energy dissipation. Indeed, Landauer’s Principle demonstrates that erasing information incurs a thermodynamic cost, linking computation to entropy generation and irreversibility^61^. In that line, artificial computational systems, including classical and reversible computers, dissipate energy as they transition between states, further emphasizing that computation is fundamentally a non-equilibrium process^63^. Therefore, the brain dynamics closer to equilibrium under psychedelics can be interpreted from the point of view of computation and interference with the brain’s “modelling engine”^64–66^. This might result in probabilistic inferences about the internal and external environment being temporarily undermined, leading to a breaking of the current models in favor of greater modelling unpredictability. At the neurophenomenological level, this might reflect the reported experiences of ego-dissolution (losing the model of the self), perceptual alterations, and altered sense of meaning^8,30,67^. This view is also consistent with the theory of relaxed beliefs under psychedelics (REBUS), positing that increased complexity in the psychedelic state is associated with reduced top-down information flow, enabling the revision and de-weighting of overweighted priors.

Living systems, guided by non-equilibrium thermodynamics, continuously consume energy to maintain ordered structures and perform computations such as gene regulation and neural signaling^68^. This implies that the non-equilibrium nature of a system is intrinsically dependent on the spatiotemporal scaling. In other words, if the system at microscopic scales reflects mechanisms that induce the breaking of the detailed balance, it is not guaranteed that the non-equilibrium dynamics are preserved at the macroscopic scale^69^. The same question has been posed in the context of brain dynamics, and, while in general, it is agreed that at the neuronal scale, the processes are out-of-equilibrium, it is less clear for macroscopic brain dynamics. Using different methodologies, recent studies have investigated and found, at the macroscopic scale, a relationship between breaking of the detailed balance with cognitive functions^21,70^, states of consciousness^19,20,71^, and neurological disorders^24^. Consequently, we also wanted to investigate how different spatial resolutions at the macroscopic level, defined by various brain parcellations, impact the non-equilibrium nature of the psychedelic experience. We found that finer parcellations exhibit a clearer breakdown of temporal asymmetry under psychedelics in comparison to coarser parcellations (**Figure 2B**). Overall, this suggests that information flow at finer scales is altered and hence, induces computational changes that predominantly occur at these scales.

### Moving closer to the equilibrium

Previous studies have indicated that reduced states of consciousness, such as sedation, general anaesthesia, and disorders of consciousness, consistently exhibit a flattening of the functional hierarchy. Huang and colleagues extensively demonstrated that these states show a contraction in the different components of spatial differentiation^10^, while Luppi and colleagues reported similar findings in anaesthetised macaques under different drugs and levels of sedation^11^. Using the thermodynamically grounded definition of hierarchy employed in this work, previous studies have also observed the flattening of the hierarchy in disorders of consciousness, deep sleep^24,72^ and movie watching^23^, exhibiting brain dynamics closer to equilibrium. Across species, this flattening has also been observed in anaesthetised and sleeping macaques^19^. These results are typically interpreted as showing a reduced repertoire of dynamics states compared to the normal wakeful state, likely indicating a reduced computational activity and less flexibility^23^.

Interestingly, the psychedelic experience has also been consistently shown to flatten the hierarchy, not only through the contraction of different components of spatial differentiation but also, as demonstrated in this work, through the flattening of the Directed Functional Hierarchy. However, unlike reduced states of consciousness, the psychedelic state is associated with more complexity, reflecting an opposite trend. This has been shown in previous studies demonstrating that the brain complexity is reduced under the effect of anaesthesia in humans^73^, rats^74^, and monkeys ^75^, as well as during sleep stages in humans^76^ and rats^77^, while brain complexity is increased under psychedelics in humans^1,46,78–81^. This suggests that both states may induce the flattening of the hierarchy through different mechanisms. We hypothesise that understanding brain complexity in this regard might help to resolve this seeming dichotomy. In support of this hypothesis, we observed that the flattening in the Directed Functional Hierarchy is inversely related to changes in brain complexity (as measured by LZ complexity) (**Figure S4-6**). In this work, the two dimensions, DFH together with brain complexity, disentangle the reduced states of consciousness from the psychedelic state concerning the thermodynamic equilibrium. Therefore, we argue that the thermodynamic equilibrium shown by dynamics in different states of consciousness can only be understood while taking a multidimensional lens of altered states of consciousness^82,83^.

The collapse of Directed Functional Hierarchy also reflects the promising therapeutic potential of psychedelics^84–87^. In pathological states such as depression, the brain’s dynamic landscape (here defined in terms of DFH) becomes dynamically trapped in local minima. Behaviorally, this is often reflected in ruminative and low mood states of depressed patients^47,88^. Moreover, these “canalised” dynamics might relate to synaptopathy through stress-induced atrophy and/or problematic Hebbian plasticity, reinforcing an inability to escape from specific states^48,89^. In this context, psychedelics—through their action on serotonin receptors—collapse or flatten functional hierarchies by increasing the complexity of neural dynamics, possibly leading to more flexible and adaptable neural states while destabilising maladaptive, ‘trapped’ ones^47^. Psychedelics appear to promote a homeostatic plasticity that could aid the recalibration or reset of canalized circuitry and dynamics, aiding psychological treatment after the interventions through the promotion of an extended period of plasticity^47^. Indeed, psychotherapy has been considered an important—if not essential—part of psychedelic treatments^90^. Moreover, novel interventions such as non-invasive brain stimulation techniques have been proposed^65^.

Looking at the thermodynamic equilibrium through the aforementioned multidimensional lens enables further characterizations of brain states in health, disease, and altered states of consciousness opening up new avenues for diagnosis and treatment predictions. Importantly, our work aims to investigate the temporal asymmetry obtained through the time-shifted correlation, which is often used as a straightforward signature of deviation from equilibrium^91^. Equally, there are different approaches for capturing the breaking of the detailed balance and temporal asymmetry that can be used to investigate the reshaping of the DFH during psychedelics. For instance, Granger Causality or Mutual Information could be considered to quantify the directionality of the information flow between brain signals^92,93^ Also different approaches capturing the non-equilibrium nature of brain dynamics could be considered, such as the generative effective connectivity as proposed by Kringelbach and colleagues^70^, or violations from the fluctuation-dissipation theorem proposed by Deco and colleagues^94^.

## Conclusion

In conclusion, our findings highlight the pivotal role of DFH in characterizing the functional reorganization of the brain under psychedelics, revealing a transition toward a more flexible and accessible cortical hierarchy. These results underscore the mechanistic link between DFH reduction and increased global brain complexity, suggesting a distinct pathway through which psychedelics bring the brain closer to thermodynamic equilibrium, potentially aiding homeostatic plasticity.

## Methods

In this section, we provide all the details of the empirical neuroimaging data for all three datasets used in this study.

### Participants

#### DMT

The full description of the dataset can be accessed in^8,95^. Twenty-five participants were considered for the experiment in a single-blind, placebo-controlled, and counter-balanced design. To be considered the participants were required to be older than 18 years of age, lack experience with a psychedelic, not have a previous negative response to a psychedelic, and/or to currently suffer from or have a history of psychiatric or physical illness. Twenty participants completed the entire study (average age = 33.5 years, SD = 7.9, 7 female). Seventeen participants were used in the analysis after further removal of three participants due to excessive motion during the 8 minutes DMT recording (more than 15% of volumes scrubbed with framewise displacement (FD) of 0.4 mm).

#### LSD

The full description of the dataset can be accessed in^30^. Twenty healthy participants were considered for the experiment (four females, average age = 30.9 ± 7.8 years) in a balanced order, within-participants design. To be considered the participants were required to be older than 21 years of age, not pregnant, not have a personal history of diagnosed psychiatric illness, not have an immediate family history of a psychotic disorder, not have previous experience with other classical psychedelic drugs, to not have any psychedelic drug use within six weeks of the first scanning day, pregnancy, to not have problematic alcohol use (i.e., >40 units consumed per week), or a medically significant condition. Fifteen subjects were considered for the current analysis. Fifteen participants were used in the analysis after the further removal of one participant due to scanner anxiety and four participants due to excessive motion during the recording (more than 15% of volumes scrubbed with framewise displacement (FD) of 0.5 mm).

#### Psilocybin

The full description of the dataset can be accessed in^29^. Fifteen participants were considered for the experiment in a counter-balanced design. To be considered the participants were required to be older than twenty-one years of age, not be pregnant, not to have a history of psychiatric disorders, not to have a cardiovascular disease, not to have a substance dependence, not to have a claustrophobia, not to have a blood or needle phobia, or to have an adverse response to hallucinogens. Furthermore, six participants were excluded if they exceeded 0.4 mm of the mean framewise displacement

### Experimental Paradigm

#### DMT

Every participant underwent two days of scanning, separated by two weeks, with two scanning sessions on each day. The first scan’s duration was twenty-eight minutes in an eyes-closed condition (with an eye mask). The eighth minute marked administration of the intravenous DMT or placebo (saline) (50/50, DMT/Placebo). Subjective effects assessment was carried out after the scanning. For the second session, the same protocol was carried out but with the assessment of subjective intensity scores. Furthermore, simultaneous EEG was recorded during the sessions.

#### LSD

Every participant underwent three scanning sessions. The scan duration was carried out after sixty minutes acclimatization period from the bolus injections (over 2 minutes) with LSD (75 μg in 10 ml saline). Three fMRI scans were carried out: rest 1 - eyes-closed resting-state session, rest 2 - resting-state session while listening to music, rest 3 - eyes-closed resting-state session. After each scan, a Visual Analog Scale (VAS) rating was carried out for each participant. We report results for the average of resting state one and resting state three scans.

#### Psilocybin

Every participant underwent two days of scanning, separated by a minimum of one week. The scan duration lasted twelve minutes with six minutes of recording followed by intravenous administration of either psilocybin (2 mg dissolved in 10 mL of saline, 60-s injection) or a placebo (10 mL of saline, 60-s injection) in an eyes-closed condition. After each scan, a Visual Analog Scale (VAS) rating was carried out for each participant.

### Acquisition Parameters

#### DMT

A 3T scanner (Siemens Magnetom Verio syngo MR 12 with compatibility for EEG recording) was used for the experiment with T2-weighted echo planar sequence and T1-weighted structural scans^8^. Relevant to the analysis in this paper, the scanning parameters were: TR/TE = 2000ms/30ms, acquisition time = 28.06 minutes, flip angle = 80o, voxel size = 3×3×3 mm3 and 35 slices with 0 mm interslice distance.

#### LSD

A 3T scanner (GE HDx system) was used for the experiment^30^. High-resolution anatomical images were acquired with 3D fast spoiled gradient echo scans in an axial orientation, with a field of view = 256 × 256 × 192 and matrix = 256 × 256 × 129 to yield 1-mm isotropic voxel resolution. TR/TE = 7.9/3.0 ms; inversion time = 450 ms; flip angle = 20. BOLD-weighted fMRI data were acquired using a gradient-echo planar imaging sequence, TR/TE = 2,000/35 ms, field of view = 220 mm, 64 × 64 acquisition matrix, parallel acceleration factor = 2, 90 flip angles. Thirty-five oblique axial slices were acquired interleaved, each 3.4 mm thick with zero slice gap (3.4-mm isotropic voxels). The precise length of each of the scans was 7 min 20 s.

#### Psilocybin

Identical acquisition was carried out for the psilocybin dataset^29^ with the following exceptions: fMRI data were acquired at TR/TE = 3,000/35 ms, field of view = 192 mm. Thirty-three oblique axial slices were acquired interleaved, each 3 mm thick with zero slice gap (3 × 3 × 3 mm voxels). The length of the scan was 97 TRs.

### fMRI Pre-processing

For fMRI pre-processing, a pipeline developed for Psilocybin^29^ and further applied to LSD^30^ and DMT^8^ experiments was used. Briefly, the following steps were applied 1) despiking, 2) slice-timing correction, 3) motion correction, 4) brain extraction, 5) rigid body registration to structural scans, 6) non-linear registration to 2mm MNI brain, 7) motion-correction scrubbing, 8) spatial-smoothing (FWHM) of 6 mm, 9) bandpass filtering into the frequency range 0.01-0.08 Hz, 10) linear and quadratic detrending, 11) regression of 9 nuisance regressors (3 translations, 3 rotations, and 3 anatomical signals). Lastly, the timeseries were parcellated into the Schaefer parcellation of different scales (100, 200, 300, 400, 500, and 1000) in the MNI space.

### Temporal asymmetry

We consider two signals, *x_i_*(*t*) and *x_i_*(*t*). First, we define the (directed) temporal (or thermodynamic) asymmetry matrix,

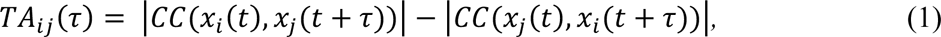

where *CC*(·,·) denotes the Pearson correlation coefficient between the two timeseries. This expression compares how well the present state of *x_i_*(*t*) predicts the future state *x_i_*(*t* + τ) versus how well the present state of *x_i_*(*t*) predicts the future state *x_i_*(*t* + τ). If *TA_ij_*(τ) > 0, it suggests that *x_i_*(*t*) provides more predictive information about *x_i_*(*t* + τ) than vice versa, indicating a forward direction of information flow from *x_i_* to *x_i_*. If *TA_ij_*(τ) < 0, it suggests the opposite direction, and if *TA_ij_*(τ) ≈ 0, no clear directional bias is discerned. By using the symmetry property of circular cross-correlations, we can rewrite *TA_i_*(τ) as

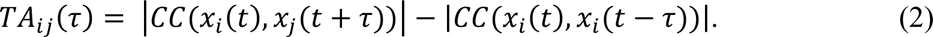

This alternate form shows that *TA_ij_*(τ) directly measures how the correlation changes when we shift *x_j_* forward in time by τ seconds versus shifting it backward by τ seconds, relative to *x_i_*(*t*).

We now define the magnitude of TA as the square value of *TA_ij_*(τ).

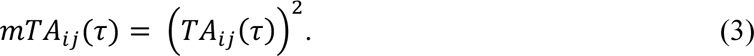

If the signals are perfectly time-symmetric, shifting one signal forward or backward by τ seconds yields the same correlation with the other, making *MTA_ij_*(τ) = 0. A nonzero value indicates that the correlation depends on the direction of the time shift, revealing a “break” in temporal symmetry. In other words, *MTA_ij_*(τ) quantifies how differently the two signals correlate when looking forward versus backward in time, helping to identify processes where the flow of information or causal influence is not symmetric with respect to time.

Finally, we quantify the (scalar) magnitude of this asymmetry as the Directed Functional Hierarchy (DFH), which is obtained as the ratio between the mean and standard deviations of all entries in the *mTA_ij_*(τ):

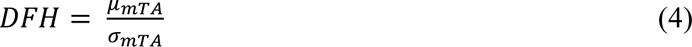

Where μ*_A_* and σ*_A_* are the mean and standard deviation of the magnitude of temporal asymmetry matrix entries, respectively. For the global DFH, the Fano factor is obtained for all regions of the magnitude of temporal asymmetry matrix. For the network-based analysis of the DFH, the matrix entries of the magnitude of temporal asymmetry matrix related to a specific resting-state network are considered.

### Components of functional differentiation of DFH

Functional components of temporal asymmetry of information flow were calculated using the BrainSpace toolbox https://github.com/MICA-MNI/BrainSpace^34^ as implemented in MATLAB. The input is the magnitude of temporal asymmetry matrix *I_ij_* obtained for each condition in each dataset represented by a *NxN* matrices, where *N* is the number of brain regions within the parcellation.

We initialized the algorithm with “*normalized angle*” kernel, “*principal component analysis*” approach and “*Procrustes analysis*” alignment. This resulted in the following steps of the algorithm: First, each row of the temporal asymmetry matrix *I_ij_* was thresholded to retain the top 10% of connections, reflecting the strongest magnitudes of temporal asymmetry for each region. Subsequently, the cosine similarity was computed between the connectivity profiles of different regions to construct a similarity matrix, denoted as *S*. Cosine similarity quantifies the similarity between two vectors by measuring the cosine of the angle between them, effectively capturing the similarity in their connectivity patterns. The cosine similarity between two vectors *u̅* and *v̅* is defined as:

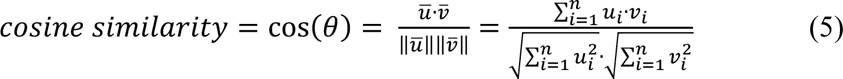

where *u̅* · *v̅* denotes the dot product of *u̅* and *v̅*, and *u̅* and *v̅* represent their magnitudes. This similarity matrix *S* is square and symmetric, encoding the similarity between regions in terms of their temporal asymmetry patterns. Given that a symmetric matrix *S* can be decomposed as *S* = *S*Σ*S^T^*, where *U* is an orthogonal matrix of eigenvectors and Σ is a diagonal matrix of eigenvalues, we employed principal component analysis (PCA) for dimensionality reduction. PCA is a linear technique that transforms the data into a low-dimensional space defined by orthogonal components that maximize variance. For each dataset, gradient analysis was performed on the averaged magnitude of the temporal asymmetry matrix. Individual subjects’ matrices were then projected onto the principal components using Procrustes alignment, as facilitated by the BrainSpace toolbox^34^.

This dimensionality reduction technique aims to uncover a meaningful low-dimensional representation of the high-dimensional original space (in this case, 400 brain regions from the Schaefer 400 parcellation). In this reduced space, the proximity of two points indicates a similar pattern of temporal asymmetry between two regions, whereas greater separation signifies differing patterns. Each gradient represents an axis of covariance in the similarity between regional patterns of temporal asymmetry. Regions with strong loadings on a particular gradient anchor that gradient, while regions with minimal loadings do not exhibit the associated pattern of variation. Notably, for each condition and subject, these anchor points may differ. The greater the disparity between minimum and maximum anchor points, the more pronounced the pattern of temporal asymmetry along the components.

To assess these variations, we quantified how the maximum and minimum values along the first two components change in the different conditions by measuring the distance between the two extremes for each components named “components 1 and 2 surface”.

### Lempel-Ziv Complexity

Consistent with Median and coleagues^80^, we first demeaned (mean value set to 0) and binarized the regional timeseries. The binarization was done by converting each timepoint into “1” if above the mean and “0” if below the mean. To derive Lempel-Ziv Complexity (LZ complexity), we used the following calculation

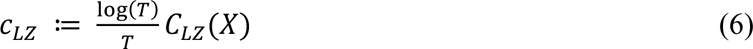

where LZ complexity can be interpreted as an efficient estimator of the entropy rate of X. In other words, the entropy rate quantifies the amount of new information, in bits, that each incoming data sample contributes, and thus carries information on how hard it is to predict the next input in the incoming data^80^. The global LZ complexity was calculated by averaging all the regional values. This allowed us to derive a simple and principled measure of brain complexity based on LZ complexity. While LZ complexity is a robust measure, it is to be noted that there are many other measures of brain complexity reflecting various aspects of spatio-temporal brain dynamics^45,46,81,96–98^. Here we wanted to show the relationship between reduced Directed Functional Hierarchy and brain complexity, and therefore chose the most traditional measure.

## Author Contributions

J.V., Y.S-P., G.D. and M.L.K. conceived the analysis. R.L.C-H designed the original study. J.V., Y.S-P., G.D. and M.L.K. designed and developed the methodologies. J.V. analysed the data. C.T. and L.R. acquired and provided the data. J.V., Y.S-P., R.L.C-H, E.T., E.L-S., E.G-G., G.R. revised the paper and provided critical feedback.

