## Supplementary material for "Brain dynamics of classical psychedelics show paradoxical hierarchical flattening with increased complexity": Brain dynamics of classical psychedelics show paradoxical hierarchical flattening with increased complexity SI - biorxiv.pdf

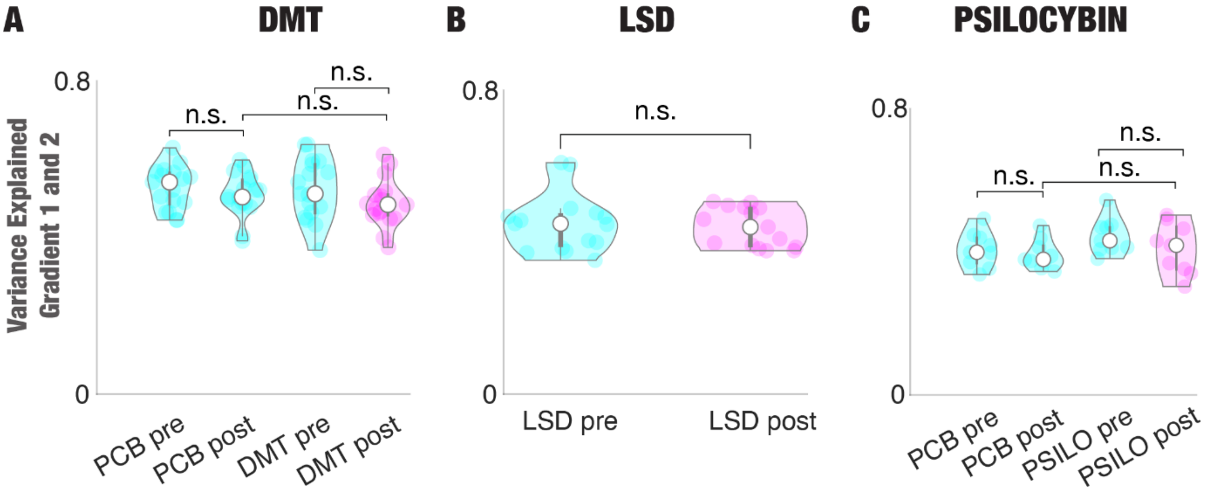


Figure S1: **The first two gradients explained the same amount of variance across conditions in the three psychedelic datasets – DMT, LSD and Psilocybin.** The amount of variance explained of the first two gradients (gradient 1 and 2) for all of the conditions in the DMT, LSD and Psilocybin datasets are not statistically significant (p-value>0.05).

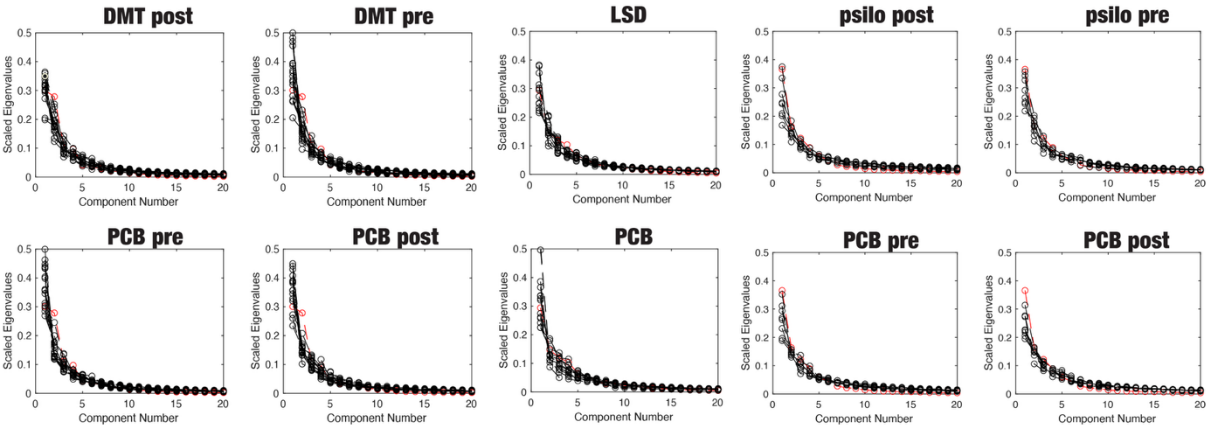


Figure S2: **Variance explained per subject and condition.** The variance explained as defined by the scaled eigenvalue for the first 20 principal components of the gradient analysis related to the magnitude of temporal asymmetry matrix for all the conditions. In red, the group average is reported and in black the individual subjects.

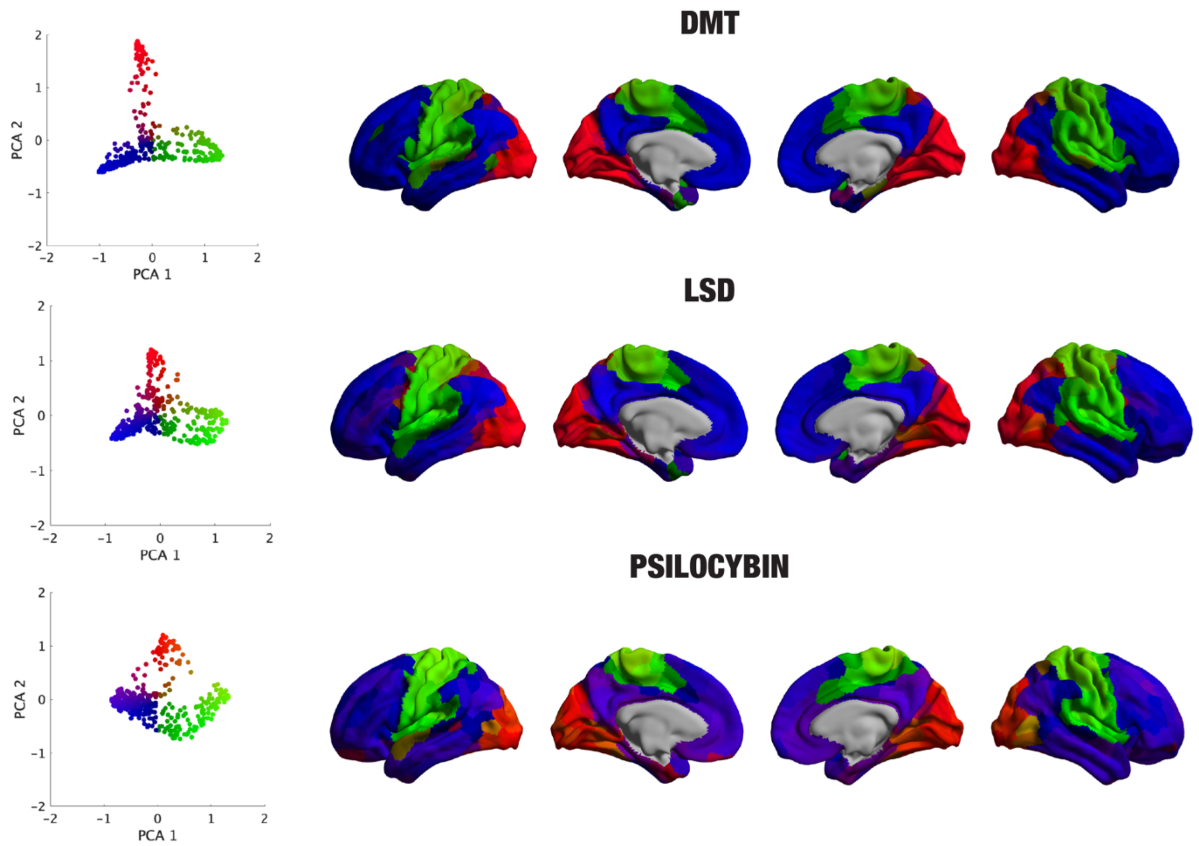


Figure S3: **2D cortical projection of the first two functional gradients of the magnitude of temporal asymmetry.** We plot the first two principal components of the magnitude of temporal asymmetry in a 2D plots and rendered on the brain surface.

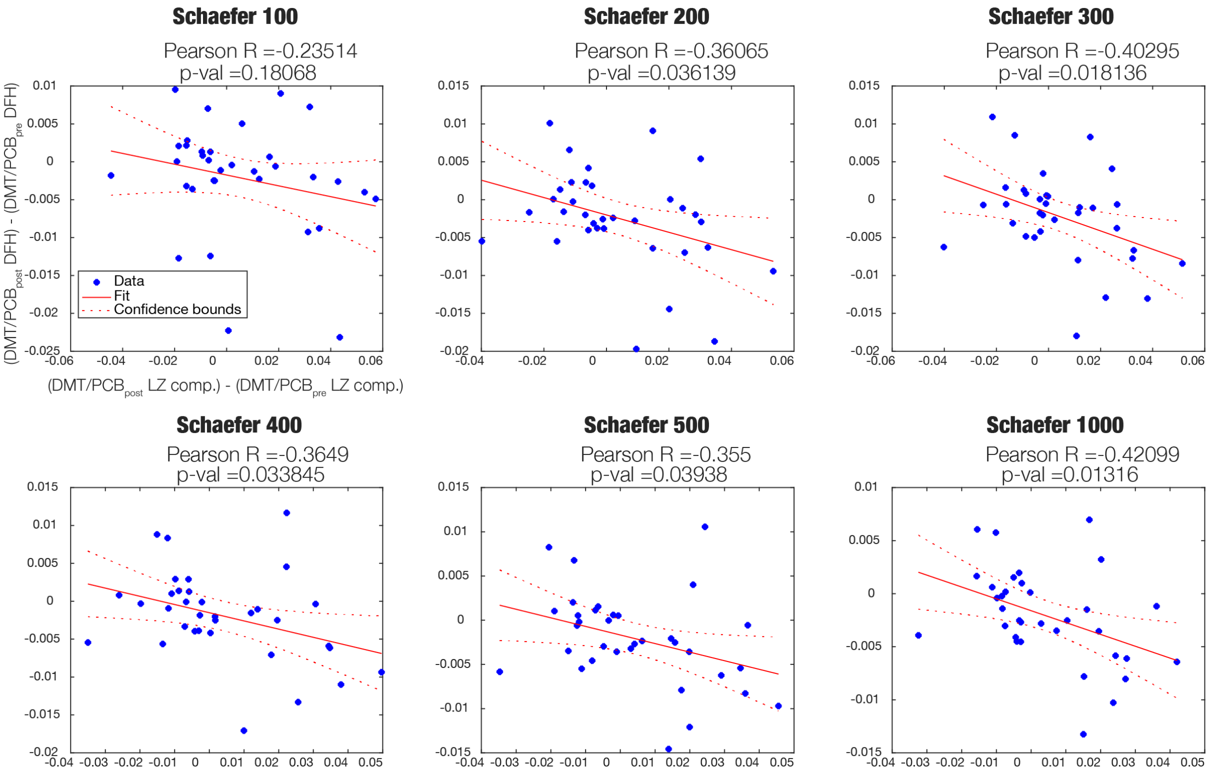


Figure S4: **The change in directed functional hierarchy negatively correlates with the change in Lempel-Ziv Complexity in the DMT dataset.** Pearson's correlation between the Directed Functional Hierarchy (DFH) and Lempel-Ziv Complexity (LZ comp.) for the DMT dataset across different Schaefer parcellations (Schaefer 100: Pearson's r = -0.24 , p-value = 0.18, Schaefer 200: Pearson's r = -0.36 , p-value = 0.036, Schaefer 300: Pearson's r = -0.40 , p-value = 0.18, Schaefer 400: Pearson's r = -0.36 , p-value = 0.034, Schaefer 500: Pearson's r = -0.36 , p-value = 0.04, Schaefer 1000: Pearson's r = -0.42 , p-value = 0.01).

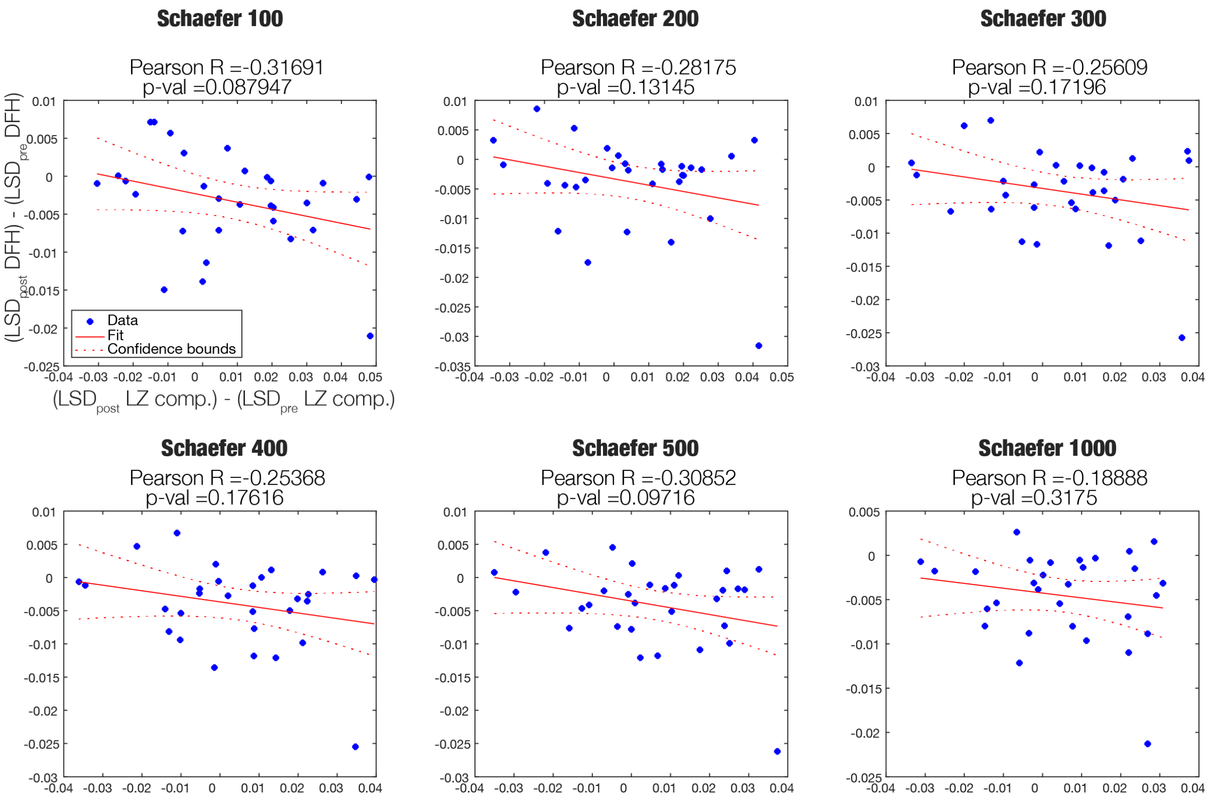


Figure S5: **The change in directed functional hierarchy negatively correlates with the change in Lempel-Ziv Complexity in the LSD dataset.** Pearson's correlation between the Directed Functional Hierarchy (DFH) and Lempel-Ziv Complexity (LZ comp.) for the LSD dataset across different Schaefer parcellations (Schaefer 100: Pearson's r = -0.31 , p-value = 0.09, Schaefer 200: Pearson's r = -0.28 , p-value = 0.13, Schaefer 300: Pearson's r = -0.26 , p-value = 0.17, Schaefer 400: Pearson's r = -0.25 , p-value = 0.18, Schaefer 500: Pearson's r = -0.31 , p-value = 0.10, Schaefer 1000: Pearson's r = -0.19 , p-value = 0.32).

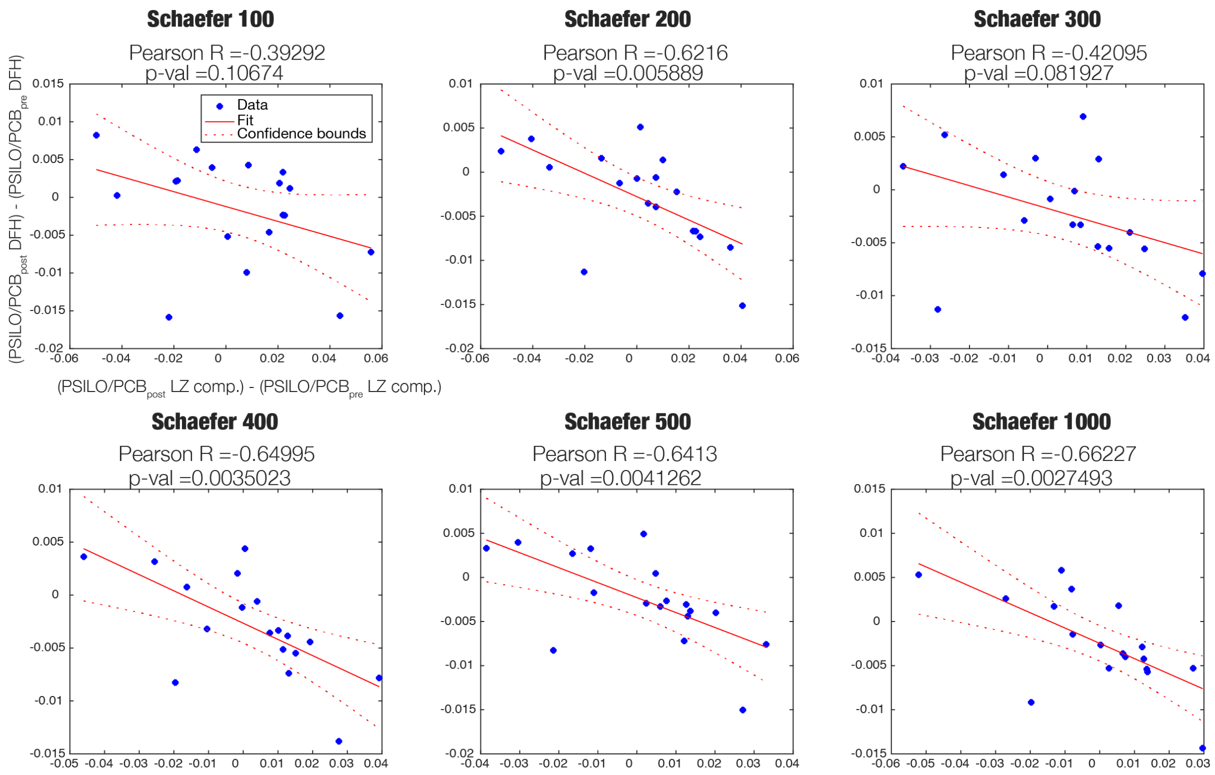


Figure S6: **The change in directed functional hierarchy negatively correlates with the change in Lempel-Ziv Complexity in the Psilocybin dataset.** Pearson's correlation between the Directed Functional Hierarchy (DFH) and Lempel-Ziv Complexity (LZ comp.) for the Psilocybin dataset across different Schaefer parcellations (Schaefer 100: Pearson's r = -0.39 , p-value = 0.107, Schaefer 200: Pearson's r = -0.62 , p-value = 0.006, Schaefer 300: Pearson's r = -0.42 , p-value = 0.082, Schaefer 400: Pearson's r = -0.65 , p-value = 0.004, Schaefer 500: Pearson's r = -0.64 , p-value = 0.004, Schaefer 1000: Pearson's r = -0.66 , p-value = 0.003).

**Table S1: Uncorrected p-values for the Network-level analysis.** Summary of the statistical tests for between-network directed functional hierarchy for each of the datasets and conditions.

**A**

**PCB post vs. PCB pre**

| | VIS | SMT | DAN | VAN | LBC | FPA | DMN |
|---|---|---|---|---|---|---|---|
| VIS | 0 | 0.38 | 0.49 | 0.53 | 0.28 | 0.19 | 0.74 |
| SMT | 0 | 0 | 0.74 | 0.27 | 0.075 | 0.92 | 0.13 |
| DAN | 0 | 0 | 0 | 0.14 | 0.064 | 0.3 | 0.88 |
| VAN | 0 | 0 | 0 | 0 | 0.11 | 0.61 | 0.057 |
| LBC | 0 | 0 | 0 | 0 | 0 | 0.75 | 0.015 |
| FPA | 0 | 0 | 0 | 0 | 0 | 0 | 0.7 |
| DMN | 0 | 0 | 0 | 0 | 0 | 0 | 0 |

**DMT post vs. DMT pre**

| | VIS | SMT | DAN | VAN | LBC | FPA | DMN |
|---|---|---|---|---|---|---|---|
| VIS | 0 | 0.0039 | 0.029 | 0.99 | 0.42 | 0.46 | 0.48 |
| SMT | 0 | 0 | 0.014 | 0.021 | 0.03 | 0.94 | 0.79 |
| DAN | 0 | 0 | 0 | 0.085 | 1 | 0.018 | 0.099 |
| VAN | 0 | 0 | 0 | 0 | 0.53 | 0.0042 | 0.21 |
| LBC | 0 | 0 | 0 | 0 | 0 | 0.87 | 0.066 |
| FPA | 0 | 0 | 0 | 0 | 0 | 0 | 0.0087 |
| DMN | 0 | 0 | 0 | 0 | 0 | 0 | 0 |

**B**

**PCB post vs. PCB pre**

| | VIS | SMT | DAN | VAN | LBC | FPA | DMN |
|---|---|---|---|---|---|---|---|
| VIS | 0 | 0.58 | 0.35 | 0.056 | 0.73 | 0.17 | 0.6 |
| SMT | 0 | 0 | 0.27 | 0.32 | 0.74 | 0.88 | 0.39 |
| DAN | 0 | 0 | 0 | 0.85 | 0.31 | 0.59 | 0.8 |
| VAN | 0 | 0 | 0 | 0 | 0.51 | 0.42 | 0.5 |
| LBC | 0 | 0 | 0 | 0 | 0 | 0.03 | 0.49 |
| FPA | 0 | 0 | 0 | 0 | 0 | 0 | 0.16 |
| DMN | 0 | 0 | 0 | 0 | 0 | 0 | 0 |

**PSILO post vs. PSILO pre**

| | VIS | SMT | DAN | VAN | LBC | FPA | DMN |
|---|---|---|---|---|---|---|---|
| VIS | 0 | 0.099 | 0.19 | 0.58 | 0.87 | 0.63 | 0.012 |
| SMT | 0 | 0 | 0.21 | 0.93 | 0.38 | 0.89 | 0.34 |
| DAN | 0 | 0 | 0 | 0.22 | 0.11 | 0.028 | 0.19 |
| VAN | 0 | 0 | 0 | 0 | 0.01 | 0.63 | 0.37 |
| LBC | 0 | 0 | 0 | 0 | 0 | 0.15 | 0.078 |
| FPA | 0 | 0 | 0 | 0 | 0 | 0 | 0.032 |
| DMN | 0 | 0 | 0 | 0 | 0 | 0 | 0 |

**C**

**LSD post vs. LSD pre**

| | VIS | SMT | DAN | VAN | LBC | FPA | DMN |
|---|---|---|---|---|---|---|---|
| VIS | 0 | 0.031 | 0.027 | 0.02 | 0.069 | 0.033 | 0.0055 |
| SMT | 0 | 0 | 0.027 | 0.061 | 0.017 | 0.69 | 0.45 |
| DAN | 0 | 0 | 0 | 0.053 | 0.23 | 0.13 | 0.78 |
| VAN | 0 | 0 | 0 | 0 | 0.11 | 0.014 | 0.37 |
| LBC | 0 | 0 | 0 | 0 | 0 | 0.0065 | 0.0035 |
| FPA | 0 | 0 | 0 | 0 | 0 | 0 | 0.0023 |
| DMN | 0 | 0 | 0 | 0 | 0 | 0 | 0 |
